# Machine-Learning-Assisted Exploration of High Entropy-Atom Nanozyme for Anti-Tumor Immunotherapy by Enhancing Enzyme Activity and Disrupting Dual Energy Metabolism

**DOI:** 10.1101/2025.07.24.666688

**Authors:** Mingyu Jiang, Zhiling Zhu

**Affiliations:** College of Materials Science and Engineering, Qingdao University of Science and Technology, Qingdao 266042, Shandong, P. R. China

**Keywords:** High Entropy-Atom Nanozyme, energy metabolism, machine learning, cancer immunotherapy

## Abstract

Despite its potential in cancer therapy, single-atom nanozyme (SAzyme) faces challenges like low atomic loading and rapid cancer metabolism. Here, a high-entropy atom nanozyme (HEAzyme), PtNiBiSnSb-anti-CD36, is designed for efficient anti-tumor immunotherapy. The PtNiBiSnSb HEAzyme, incorporating five SAzyme, demonstrates enhanced peroxidase (POD)-like activity due to its abundant active sites, showing a 7.2-fold increase in catalytic efficiency compared to the structurally analogous PtBi nanozyme. Further integrating of density functional theory calculations and machine learning analysis reveals that the main reason is due to the addition of Ni, Sn, and Sb, which reduced the reaction energy barrier and changed the adsorption energy of surface hydroxyl groups. Moreover, PtNiBiSnSb-anti-CD36 not only inhibits normal metabolic processes by depleting nicotinamide adenine dinucleotide (NADH) through its enhanced NADH-POD-like activity, but also suppresses lipid metabolism by blocking CD36-mediated uptake. The dual metabolic inhibition facilitates the mild photothermal therapy induced by PtNiBiSnSb. In this process, the anti-tumor immune response is amplified by inducing immunogenic cell death and inhibiting the growth of immunosuppressive cells through blocking fatty acid uptake. The proposal of HEAzyme for cancer therapy provides a novel approach to enhance the catalytic activity of SAzymes and deepens the theoretical understanding of HEAzyme in the POD-like reaction process.

## 1. Introduction

Currently, breast cancer has become the leading cancer by incidence worldwide.^[1]^ Unlike normal cells, cancer cells exhibit significant metabolic alterations, including an abnormal increase in glucose and glutamine demand, as well as changes in lipid metabolism such as increased lipogenesis and fatty acid uptake.^[2]^ For breast cancer cells, lipid metabolism may be particularly crucial.^[3]^ Breast cancer cells are often surrounded by a large number of adipocytes, creating a fatty acid-rich microenvironment that can act as an external growth stimulus.^[4]^ Additionally, these lipid metabolic products can aberrantly accumulate in immunosuppressive cells, such as M2 tumor-associated macrophages (TAMs), myeloid derived suppressor cells (MDSCs), and regulatory T (Treg) cells, which are characterized by high expression of the CD36 receptor, mediating lipid uptake as a primary energy source.^[5]^ Therefore, blocking lipid uptake can inhibit the function of immunosuppressive cells, help restore immune surveillance in the tumor microenvironment (TME), and potentially disrupt cancer progression and improve treatment outcomes. Several studies have reported that inhibiting lipid metabolism can enhance cancer treatment.^[6]^ For example, Song et al. developed a hydrogel capable of simultaneously delivering hollow mesoporous CuS nanoparticles and the CD36 inhibitor sulfosuccinimidyl oleate to inhibit lipid metabolism.^[7]^ Yang et al. loaded the fatty acid synthase inhibitor cerulenin into an Ir-N5 SAzyme carrier for fatty acid synthesis inhibition.^[8]^ Although these materials effectively inhibit lipid metabolism, they lack specificity, posing off-target risks and insufficient catalytic activity. Therefore, it is essential to design a nanomaterial with high catalytic capacity that can specifically inhibit CD36-mediated fatty acid uptake.

Among various nanomaterials, SAzymes have garnered significant attention in nanomedicine due to their superior catalytic performance.^[9]^ However, SAzymes face limitations such as low single-atom loading and excessively strong binding between transition metal sites and electron-donating intermediates, resulting in a high energy barrier for catalytic reactions.^[10]^ High-entropy intermetallic compounds offer unique advantages as they are a class of intermetallic compounds consisting of five or more constituent elements with uniform structure and homogeneous composition.^[11]^ Compared with single metal catalysts, high entropy intermetallic compounds are with rich element combinations and diverse chemical properties.^[12]^

In contrast to high-entropy alloys, which enhance catalytic performance by incorporating multiple metal elements^[13]^, high-entropy metal intermetallic compounds, referred to as HEAzymes further utilize the interactions between adjacent metal atoms to modulate the energy barriers of reaction intermediates, enhance catalytic sites, and adjust catalytic performance. This provides a promising strategy for enhancing catalytic activity in cancer treatment. Their well-defined structure also offers new perspectives for studying the microscopic mechanisms of catalytic reactions, aiding in a deeper understanding of the role of multi-metal synergistic effects in catalysis. Although high-entropy intermetallic compounds exhibit a certain degree of order compared to high-entropy alloys, the diversity of combinations of five elements still makes it difficult to perform accurate theoretical calculations, similar to the fixed structure of pure metals. Machine learning, an important branch of artificial intelligence, has garnered significant attention due to its ability to efficiently process large-scale and high-dimensional data, extracting valuable information from it.^[14]^ At present, machine learning has already been applied in the theoretical catalysis field of high-entropy alloys.^[15]^ For instance, Song et al. applied machine learning to the hydrogen evolution reaction of multi-site catalysts, revealing the relationship between geometric structure and overall activity.^[16]^ Guo et al. constructed a machine learning model to predict millions of active sites on the surface of high-entropy alloys, explaining the structure-activity relationship between element composition and oxygen reduction reaction (ORR) catalytic activity.^[17]^ However, there are currently no reports on the use of machine learning to predict the application of HEAzyme in vivo catalytic therapy.

In this study, a PtNiBiSnSb-anti-CD36 HEAzyme was designed to enhance oxidative stress and disrupt lipid metabolism, aiming to promote anti-tumor immunotherapy. The PtNiBiSnSb HEAzyme was synthesized using a simple wet-chemical method. Due to its abundant active sites, it exhibited exceptional catalytic performance in enzyme-like reactions, including peroxidase (POD)-like and nicotinamide adenine dinucleotide (NADH)-POD-like activities. Enzyme kinetics experiments revealed that the catalytic efficiency of the POD-like activity of PtNiBiSnSb HEAzyme was 7.2 times higher than that of its structurally analogous PtBi nanozyme. By further combining density functional theory (DFT) calculations with machine learning analyses, it was elucidated that the POD-like activity gradually increased due to the incorporation of Ni, Sn, and Sb, which altered the electronic configuration on the surface of the HEAzyme, thereby affecting the adsorption energy of catalytic reaction intermediates. Moreover, PtNiBiSnSb HEAzyme demonstrated broad absorption in the second near-infrared window (NIR-II) and could generate mild heat (40-43℃) under 1064 nm laser irradiation, facilitating mild-photothermal therapy (PTT) ablation of tumor cells. The PtNiBiSnSb HEAzyme was further covalently conjugated with an anti-CD36 antibody to form PtNiBiSnSb-anti-CD36. Because of rich H_2_O_2_ and NADH in the TME, PtNiBiSnSb-anti-CD36 disturbs the NADH/NAD^+^ balance by efficiently generating •OH and oxidizing NADH to NAD^+^. This process not only reduces mitochondrial oxidative respiration by inhibiting the mitochondrial electron transport chain (ETC) but also interferes with the metabolic processes of tumor cells, impairing their ability to produce adenosine triphosphate (ATP). Interestingly, PtNiBiSnSb-anti-CD36 specifically inhibits ATP production only within the TME. The decrease in ATP levels further augments mild-PTT by diminishing the production of heat shock proteins (HSPs). Additionally, PtNiBiSnSb-anti-CD36 also inhibits CD36-mediated lipid uptake, comprehensively disrupting metabolic homeostasis. Blocking lipid uptake can suppress the function of immunosuppressive cells, further enhancing the efficacy of immunotherapy induced by enzyme-like catalyzed treatment and mild-PTT (**Scheme 1**). This work provides a paradigm for machine learning-assisted exploration of HEAzyme to enhance the catalytic activity of SAzymes for cancer immunotherapy, broadening the application of HEAzyme in the biomedical field.

## 2. Results and Discussion

### 2.1 Synthesis and Characterization of HEAzymes

The PtNiBiSnSb HEAzyme was synthesized via a one-batch method (**Figure 1a**). Based on the same hexagonal-close-packed (*hcp*) structure of PtBi, PtSn, PtSb, and NiBi intermetallic compounds, Pt, Ni, Bi, Sn, and Sb elements were rationally selected to construct the PtNiBiSnSb high-entropy intermetallic (HEI). Fixing the atomic ratio of (Pt + Ni)/(Bi + Sn + Sb) to 1:1 is the key principle to achieve a single-phase *hcp* PtBi-type HEI compound. The PtNiBiSnSb HEI shares similarities peak patterns compared with PtBi and NiBi intermetallic compounds, all exhibiting a *hcp* crystalline structure. Transmission electron microscopy (TEM) images (Figure 1b) reveal a uniform dispersion of the as-obtained product with an average particle size of 14.3 nm. The high-angle annular dark-field scanning TEM (HAADF-STEM) elemental mapping images (Figure 1c) further confirm the coexistence and uniform distribution of Pt, Ni, Bi, Sn, and Sb atoms. Subsequently, the aberration-corrected HAADF-STEM image (Figure 1d) clearly indicate the fully ordered stacking features in the PtNiBiSnSb HEI. Notably, a regular arrangement of two dark spots and one bright spot at the nanoparticle edge can be distinctly observed in the [001] plane (Figure 1e), corresponding to the Bi-type columns and Pt-type columns in the *hcp* PtBi intermetallic compound, respectively. As shown in Figure 1d, the lattice spacing of the (100) plane at the PtNiBiSnSb edge is measured to be approximately 0.364 nm, which lies between the lattice spacings of the (100) planes in the PtBi (0.374 nm), PtSn (0.355 nm), and PtSb (0.358 nm) intermetallic compounds. Furthermore, a uniform lattice spacing of 0.21 nm manifests that the entire PtNiBiSnSb possesses the fully ordered *hcp* crystal structure. The atomic ratio of Pt/Ni/Bi/Sn/Sb was determined by inductively coupled plasma mass spectrometry (ICP-MS) to be 23.8/23.7/31.4/9.9/11.2, which is similar to the energy dispersive X-ray spectroscopy (EDX) results (Pt/Ni/Bi/Sn/Sb = 24.1/23.6/28.4/13.0/10.9), indicating the high quality and purity of PtNiBiSnSb nanocrystals. Additionally, XPS analyses determined the valence states of these five elements in the PtNiBiSnSb HEI. The predominant metallic state of Pt and the presence of zero-valence states in Ni, Bi, Sn, and Sb are revealed in Figure 1f-j. In detail, Ni, Bi, Sn, and Sb elements are mainly in oxidized states, due to their higher tendency to oxidize in the air atmosphere. These results confirm the successful synthesis of the PtNiBiSnSb HEI compound, in which Ni substitutes the Pt columns, and Sn and Sb substitute Bi columns.

**Figure 1.**
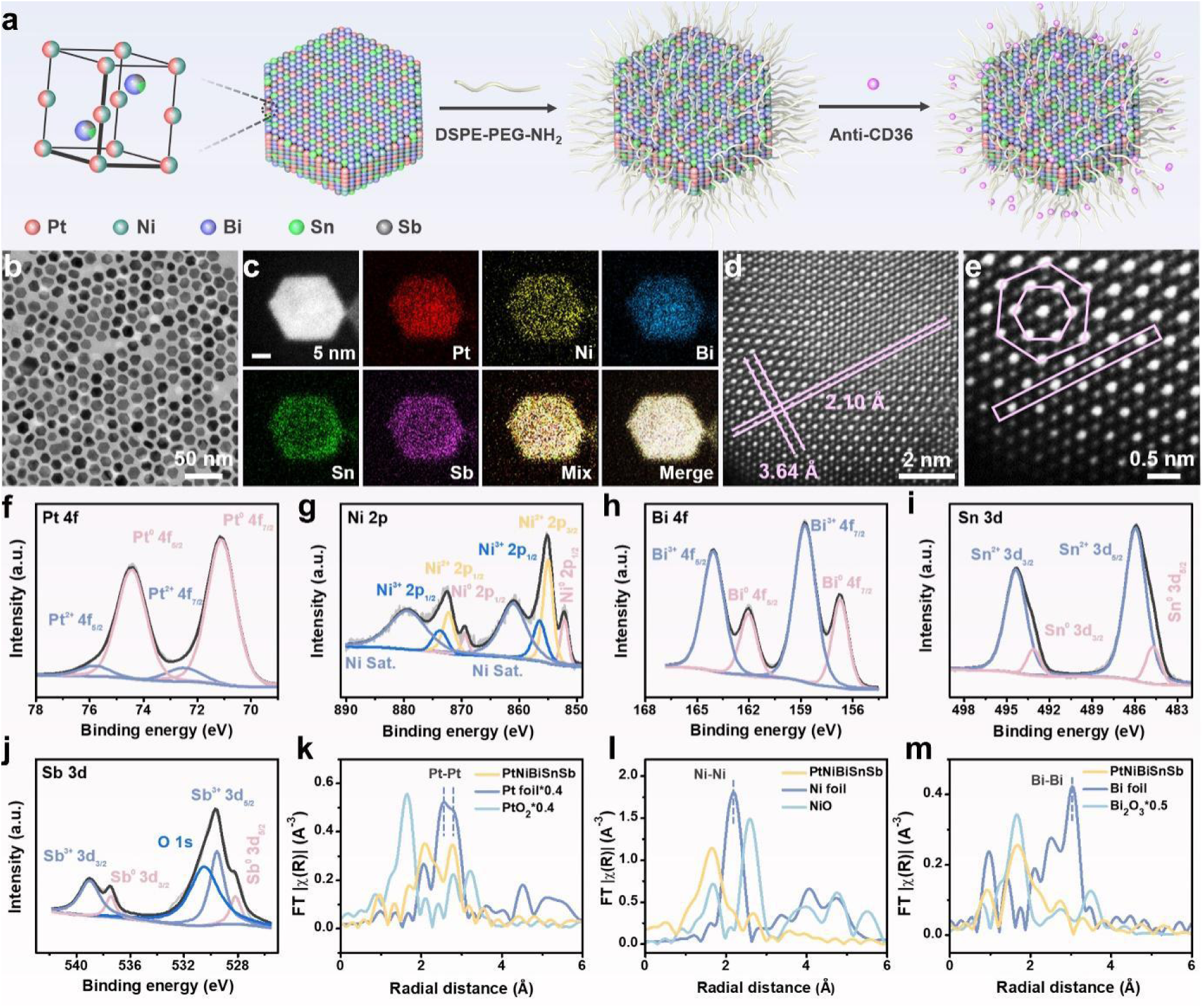
Morphology and structural characterization of PtNiBiSnSb. (a) Schematic diagram of the synthesis route of PtNiBiSnSb-anti-CD36. (b) TEM image and (c) elemental mapping images of PtNiBiSnSb. (d-e) Spherical aberration HAADF-STEM images of PtNiBiSnSb. (f-j) XPS spectra of PtNiBiSnSb. Fourier transform EXAFS spectra of (k) Pt *L*-edge, (l) Ni *K*-edge, and (m) Bi *L*-edge of PtNiBiSnSb and reference samples including Pt foil, PtO_2_, Ni foil, NiO, Bi foil, and Bi_2_O_3_.

The intrinsically isolated Pt, Ni, Bi, Sn, and Sb atoms in PtNiBiSnSb HEI were analyzed using X-ray absorption near-edge structure (XANES) and extended X-ray absorption fine structure (EXAFS) spectra. Since Pt, Ni, and Bi are the elements with higher proportions in the intermetallic compounds, while Sn and Sb substitute the Bi positions, Pt, Ni, and Bi were selected as representatives for the tests. By comparing the absorption thresholds of PtNiBiSnSb with Pt foil and PtO_2_, the average valence state of Pt is between 0 and +4, which matches with the XPS analyses. Additionally, the Fourier transform Pt-*L* edge EXAFS spectrum (Figure 1k) of PtNiBiSnSb shows the absence of Pt-Pt coordination (2.55 Å and 2.79 Å) corresponding to Pt foil and Pt-O-Pt coordination (2.80 Å and 3.22 Å) corresponding to PtO_2_, suggesting that Pt species are atomically dispersed in PtNiBiSnSb. The main scattering peaks at 1.72 Å, 2.36 Å, and 2.77 Å could be attributed to the Pt-O and multiple Pt-Ni/Bi/Sn/Sb metallic coordination in the HEI with complex coordination environment. Similarly, the XANES absorption edge of Ni *K*-edge of PtNiBiSnSb is positioned to the right of that of NiO, indicating that the oxidation state of Ni is greater than +2. Ni *K*-edge EXAFS (Figure 1l) revealed that PtNiBiSnSb has a main peak at 1.66 Å and a weak peak at 2.67 Å, attributing to Ni-O and metallic Ni-Pt/Bi/Sn/Sb coordination, respectively. There is no Ni-Ni coordination (2.18 Å) corresponding to Ni foil and Ni-O-Ni coordination (2.59 Å) corresponding to NiO, signifying the isolated Ni atoms in PtNiBiSnSb. The XANES absorption edge of Bi-*L* edge of PtNiBiSnSb is located between those of Bi foil and Bi_2_O_3_, suggesting that the oxidation state of Bi is between 0 and +3. Bi-*L* edge EXAFS (Figure 1m) showed three scattering peaks at 1.68 Å, 2.22 Å, and 2.73 Å attributing to Bi-O and metallic Bi-Pt/Ni/Sn/Sb coordination, respectively, which are different from the Bi-Bi coordination (3.04 Å) in Bi foil or the Bi-O-Bi coordination (3.49 Å) in Bi_2_O_3_, proving the isolated dispersion of Bi atoms in PtNiBiSnSb.

To ensure optimal dispersion of PtNiBiSnSb in aqueous solution, DSPE-PEG-NH_2_ was employed as a surface modifier, enabling subsequent covalent attachment of CD36 antibodies on PtNiBiSnSb via an amide reaction. Significant changes in Zeta potential and dynamic light scattering (DLS) were observed before and after covalent conjugation, shifting from +11.0 mV and 68 nm to +20.4 mV and 106 nm, respectively. PtNiBiSnSb-anti-CD36 exhibited excellent dispersibility in water, phosphate buffered saline (PBS), and Roswell Park Memorial Institute (RPMI) medium, qualifying it for in vivo experiments. Subsequently, the stability of PtNiBiSnSb-anti-CD36 in FBS-containing medium was subsequently evaluated. Both DLS and TEM analyses showed no significant changes in the nanoparticles during incubation for up to 7 days, indicating good stability. Furthermore, a hemolysis assay confirmed the excellent biocompatibility of PtNiBiSnSb-anti-CD36, even at concentrations as high as 400 μg/mL.

### 2.2 Enzyme-like Catalytic Activity and Photothermal Capacity of PtNiBiSnSb HEAzymes

Based on the abundant active sites in HEAzymes, nanozyme catalytic activities were comprehensively investigated. The highly toxic reactive oxygen species (ROS) can induce DNA damage and lipid peroxidation, exhibiting significant anti-tumor effects. Therefore, the ability of PtNiBiSnSb HEAzymes to generate ROS was studied. The POD-like activity was verified using 3,3’,5,5’-tetramethylbenzidine (TMB), which can be oxidized by hydroxyl radicals (•OH) to form blue oxidized TMB (**Figure 2a**). The absorbance peak of TMB at 652 nm gradually increased over time, reaching its maximum value within 6 min. This illustrates that the PtNiBiSnSb possesses POD-like activity and can efficiently convert H_2_O_2_ into •OH. To prove that the PtNiBiSnSb HEAzyme exhibits higher catalytic activity due to its numerous active sites, the POD-like activities of PtBi, PtBiSn, PtBiSnSb, and PtNiBiSnSb were systematically compared. First, their ability to oxidize TMB was tested under the same conditions. As shown in Figure 2b, the POD-like activity followed the trend PtNiBiSnSb > PtBiSnSb > PtBiSn > PtBi. To further quantify the POD-like activity, steady-state kinetic analysis was performed using different H_2_O_2_ concentrations. The maximum reaction velocity (V_max_) and Michaelis–Menten constant (K_m_) were obtained from the Michaelis–Menten fitting curve and Lineweaver-Burk plot (Figure 2c-j). PtNiBiSnSb exhibited the highest V_max_ (1.448×10^-6^ M s^-1^), which is 2.64, 1.48, and 1.39 times that of PtBi, PtBiSn, and PtBiSnSb, respectively. Simultaneously, PtNiBiSnSb also exhibited the lowest K_m_ (0.31 mM), implying a higher affinity between PtNiBiSnSb and the substrate. Based on the K_m_ and V_max_ values, PtNiBiSnSb exhibited superior POD-like activity. Moreover, the catalytic efficiency was calculated from the enzymatic parameters, revealing that PtNiBiSnSb has the highest K_cat_/K_m_ (18.58 mM⁻¹ s⁻¹), whereas PtBi, PtBiSn, and PtBiSnSb show values of only 2.59, 8.65, and 14.56 mM⁻¹ s⁻¹, respectively.

**Figure 2.**
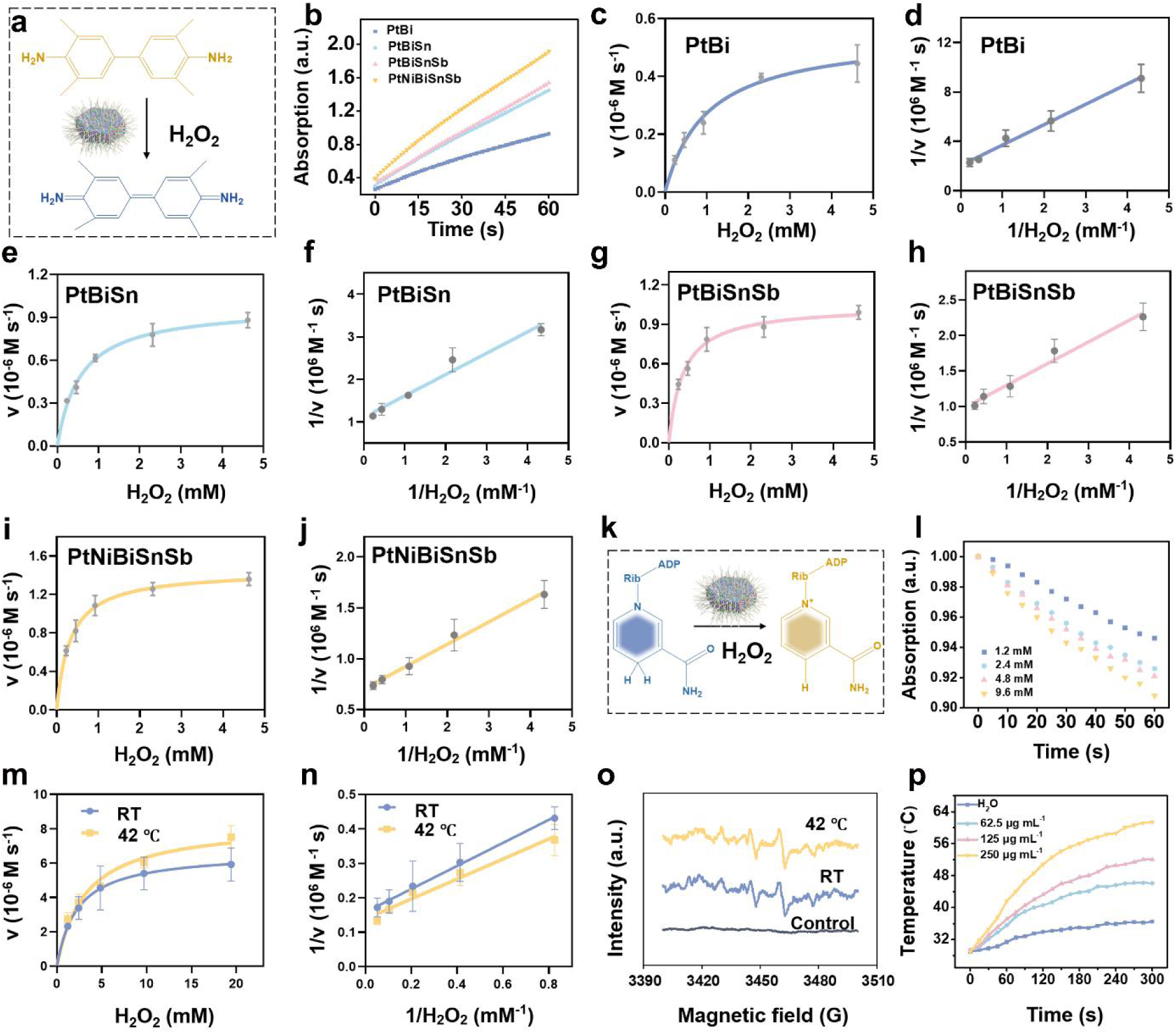
Enzyme-like activity and photothermal performance characterization of PtNiBiSnSb. (a) Schematic illustration for the reaction of TMB, which can be used for in vitro measurement of POD-like activity. (b) Time-dependent TMB absorbance changes. Michaelis−Menten kinetic analysis (c, e, g, i) and Lineweaver-Burk plotting (d, f, h, j) of POD-like activity with H_2_O_2_ as substrate, respectively. Data presented as mean ± S.D. (n = 3). (k) Schematic illustration for the reaction of NADH, which can be used for in vitro measurement of NADH-POD-like activity. (l) Time-dependent NADH absorbance changes. Michaelis−Menten kinetic analysis (m) and Lineweaver--Burk plotting (n) of NADH-POD-like activity with H_2_O_2_ as substrate, respectively. (o) ESR spectra of different groups for the detection of ·OH radical. (p) Temperature changes of PtNiBiSnSb at different concentrations under 1064 nm laser irradiation (0.5 W cm^−2^, 5 min).

NADH is an important coenzyme factor involved in cellular energy metabolism, characterized by a distinct absorption peak at 340 nm (Figure 2k). Given that PtNiBiSnSb exhibits high POD-like activity, NADH-POD-like activity was further investigated. As shown in the Figure 2l, after co-incubating PtNiBiSnSb with H_2_O_2_ and NADH, the characteristic absorption peak of NADH at 340 nm gradually decreased, indicating that PtNiBiSnSb has NADH-POD-like activity and can oxidize NADH to NAD^+^. Increasing the temperature can accelerate the reaction, enhance the binding efficiency between the nanozyme and the substrate, and promote its catalytic activity. Afterwards, the enzyme kinetics of NADH-POD-like were tested by measuring the rate of NADH consumption at room temperature and at 42°C under varying H_2_O_2_ concentrations (Figure 2m,n). The kinetic processes followed the typical Michaelis-Menten equation. At room temperature the K_m_ and V_max_ values were 2.27 mM and 6.64×10^-7^ M s^-1^, respectively, whereas at 42°C the corresponding values were 2.00 mM and 8.16×10^-7^ M s^-1^, which was 1.23 times and 0.88 times higher than the former, signifying that hyperthermic conditions are favorable for the POD-like activity of the nanozyme. Next, the generation of •OH was confirmed through electron spin resonance (ESR) experiments, which also validated that the ability to produce ROS was enhanced under mild photothermal conditions (Figure 2o). Further verification is performed to determine whether PtNiBiSnSb HEAzymes possess heat-generating capabilities to enhance nanozyme activity. The PtNiBiSnSb HEAzymes exhibit strong absorption in the NIR-II region, with absorption increasing as the material concentration increases. According to the Lambert-Beer law, the extinction coefficient of PtNiBiSnSb at 1064 nm was calculated to be 2.18 L g^-1^ cm^-1^. Subsequently, the photothermal properties of the PtNiBiSnSb under 1064 nm laser irradiation were studied using a thermal imaging camera. PtNiBiSnSb exhibits a laser power density-dependent temperature increase under 1064 nm laser irradiation. Next, the heat generation capacity of different concentrations of the PtNiBiSnSb under the same power density was measured. As shown in Figures 2p, 250 μg mL^-1^ of PtNiBiSnSb increased the temperature by 32.4℃ within 5 min under 1064 nm laser irradiation at 0.5 W cm^-2^, while water only increased by 7.6 ℃ under the same conditions. This demonstrates that PtNiBiSnSb has a strong photothermal conversion capability, with a calculated photothermal conversion efficiency of 34%. Furthermore, the photothermal stability of PtNiBiSnSb was tested. The heat generation capacity of PtNiBiSnSb did not show significant changes over four laser on-off cycles, indicating that PtNiBiSnSb has good photothermal stability. The above data collectively indicate that PtNiBiSnSb has potential for application in PTT.

### 2.3 Machine learning exploration of mechanism of PtNiBiSnSb HEAzymes

Due to the numerous microstructures on the surface of HEAzymes, applying traditional single active-site computational methods to HEAzymes is difficult and does not align with practical conditions. Gao and colleagues established a scaling relationship for POD-like activity in materials, which only requires considering a single adsorbate-specifically, hydroxyl adsorption energy-as the descriptor for catalytic activity.^[18]^ In this context, hydroxyl adsorption energy is chosen to quantitatively describe the POD-like activity of PtNiBiSnSb. However, it is clearly impractical to use DFT calculations for all sites due to the large number of active sites on the surface of PtNiBiSnSb. With its powerful data processing capabilities, machine learning holds promise for rapidly predicting hydroxyl adsorption energy with accuracy close to that of DFT calculations. The hydroxyl group tends to adsorb on the top and bridge sites of the PtNiBiSnSb surface. To accurately predict the hydroxyl adsorption energy on a large number of surface active sites of PtNiBiSnSb, atoms within 5 Å of the adsorption sites were selected to describe the atomic environment of the adsorption sites (**Figures 3a,b**;). Additionally, readily available geometric descriptors (including atomic number, atomic radius, relative atomic mass, generalized coordination number) and electronic information descriptors (electron affinity, first ionization energy, electronegativity, d-band electron count, work function) were chosen to characterize different atoms. Based on Pearson correlation analysis, five weakly correlated features were selected as inputs for the machine learning model, namely atomic radius, relative atomic mass, first ionization energy, d-band electron count, and work function (Figure 3c). Subsequently, using DFT calculation data, seven different regression models were trained to predict the hydroxyl adsorption energy on the material surface. Among the seven models, the feedforward neural network (FNN) achieved the highest R² score of 0.813 on the test set, with a root mean square error (RMSE) of 0.089 eV, and no overfitting was observed (Figures 3g,h). As shown in Figures 3d-f, the hydroxyl adsorption energy on the HEAzyme surface exhibits a continuous spectrum, with a broader distribution as the number of components increases. The hydroxyl adsorption ability of Pt active sites approaches the high-activity region (−2.6 eV) as defined by the descriptor with the doping of Ni, Sn, and Sb, indicating that the doping of elements induces the formation of a highly catalytically active surface with a unique structure, thus enhancing the POD-like activity. According to the machine learning predictions, Ni atoms in the second layer reduce the hydroxyl adsorption energy at the Pt atom adsorption sites on the surface, enhancing catalytic activity. Conversely, Pt atoms in the second layer increase the hydroxyl adsorption energy at Ni atom adsorption sites on the surface. To verify the accuracy of the machine learning results, further partial density of states (PDOS) analysis was conducted. With the increase in components, the d-band center of the Pt sites on the HEAzyme surface gradually decreases, inhibiting the excessive binding of hydroxyl groups on the HEAzyme surface (Figure 3j), thereby enhancing the material’s POD-like performance. Ni doping at Pt sites not only lowers the d-band center of the surface Pt sites but also provides additional strong hydroxyl adsorption sites, consistent with the trend of hydroxyl adsorption energy predicted by machine learning. The d-band centers of Pt and Ni sites both show an upward trend from the bulk to the surface in the PtBi intermetallic compound and HEAzyme (Figures 3k,l,m). SHapley Additive exPlanations (SHAP) analysis was used to assess the influence of the coordinating atoms and the selected features on hydroxyl adsorption energy (Figure 3i). For the top site, the nearest atom A1 to the hydroxyl group exhibited the strongest correlation, and the three equivalent atoms in the S2 region displayed similar feature importance. For the bridge site, the more distant atom A5 from the (BiSnSb) site in the S3 region exhibited the strongest feature importance. Among the selected features, the work function had the highest feature importance, consistent with the significance of electron transfer in the POD reaction.

**Figure 3.**
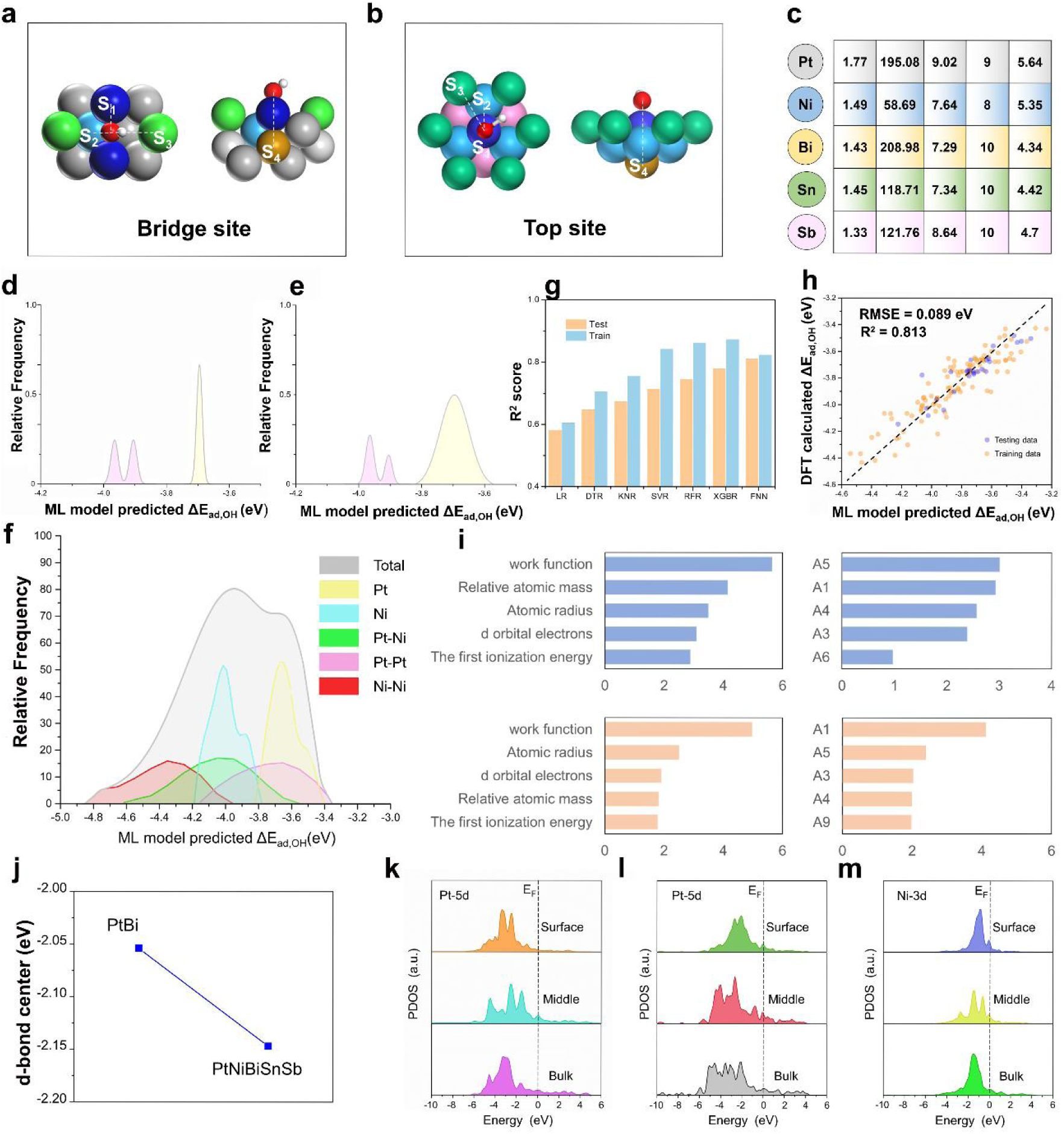
Exploring the nature of the enhanced catalytic activity of PtNiBiSnSb HEAzyme by combining DFT calculations with machine learning. Schematic diagram of bridge site (a) and top site (b). (c) Machine learning input characteristics used to describe atoms. (d,e,f) The distribution of hydroxyl adsorption energy on the surface of PtBi (d), PtBiSnSb (e) and PtNiBiSnSb (f) predicted by machine learning. (g) Performance of different machine learning models. (h) Accuracy of feedforward neural network. (i) Shapley Value Analysis of feedforward neural networks. (j) Changes in the d-band center of surface platinum atoms. (k) Site-dependent PDOSs of Pt-5d in PtBi. (l,m) Site-dependent PDOSs of Pt-5d (l) and Ni-3d (m) in PtNiBiSnSb.

### 2.3 In vitro Evaluation of Cell Apoptosis and Promotion of Immune Response

Given the excellent enzyme-like activity and photothermal conversion capabilities of the PtNiBiSnSb HEAzymes, the ability to kill tumor cells was investigated. Before demonstrating its cytotoxicity, the uptake of the nanomaterial by 4T1 tumor cells was examined. Rhodamine B was used to label thePtNiBiSnSb. As shown in **Figure 4a**, the red fluorescence intensity in the cells increased over time, confirming continuous uptake of PtNiBiSnSb by 4T1 cells, with the highest level at 4 h. Next, the potential toxicity of PtNiBiSnSb to normal cells was tested using two types of normal cells: fibroblasts (L929) cells and human umbilical vein endothelial cells (HUVECs). As shown in Figure 4b-c, PtNiBiSnSb exhibits low toxicity to these cells even at concentrations as high as 500 μg mL^-1^. This lack of toxicity may be due to the low levels of H_2_O_2_ in normal cells. The biocompatibility of PtNiBiSnSb has thus been preliminarily indicated. Subsequently, tumor cells toxicity was measured. Two breast cancer cell lines, 4T1 and MCF-7, were chosen for this test. The results showed that PtNiBiSnSb effectively killed both types of tumor cells (Figure 4d,e), which is likely due to the overexpression of H_2_O_2_ in tumor cells.^[19]^ PtNiBiSnSb can react with H_2_O_2_ to generate toxic • OH, thereby increasing oxidative stress in the TME. This hypothesis was further confirmed by the DCFH-DA assay for ROS detection.

**Figure 4.**
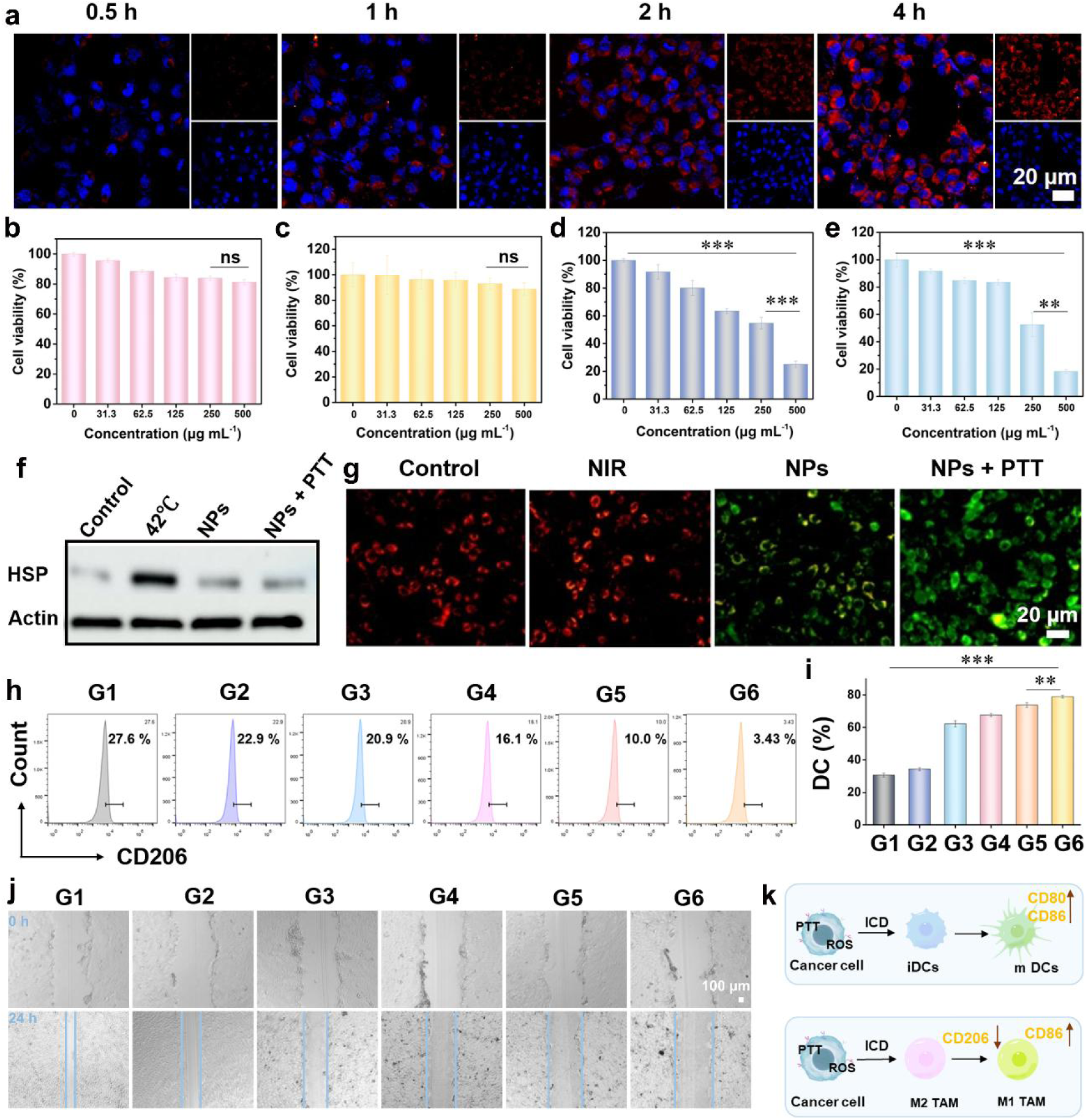
In vitro evaluation of cell apoptosis and promotion of immune response. (a) Fluorescence images of 4T1 cells incubated with PtNiBiSnSb-RhB at different time points. The cell viability of (b) HUVEC, (c) L929, (d) MCF-7, and (e) 4T1 cells after incubation with different concentrations of PtNiBiSnSb for 24 h. (f) Western blot analysis for the expressions of HSP 70 in 4T1 cells. (g) Mitochondrial membrane potential of 4T1 cells detected with JC-1 assay kit after different treatments. (h) Populations of M2 macrophages were analyzed by flow cytometry after different treatments. (i) Quantification of mature DCs by flow cytometry after different treatments. (j) Inhibition effect of PtNiBiSnSb-anti-CD36 on cell migration studied by a wound healing assay. (k) Schematic diagram of ICD promoting immune response. G1: Control, G2: anti-CD36, G3: PtNiBiSnSb, G4: PtNiBiSnSb-anti-CD36, G5: PtNiBiSnSb+1064 nm, G6: PtNiBiSnSb-anti-CD36+1064 nm. *p < 0.05, **p < 0.01, and ***p < 0.001 by Student’s two-tailed t test.

Additionally, PtNiBiSnSb demonstrates an enhanced killing effect under 1064 nm laser irradiation due to the bidirectional promotion of mild-PTT and enzyme-like catalyzed therapy. Mild-PTT enhances enzyme-like activity, thereby increasing the catalytic capability of PtNiBiSnSb. On the other hand, the •OH produced by enzyme-like catalysis consume NADH, which interferes with metabolism, leading to a decrease in ATP levels. This results in lower HSP levels (Figure 4f), thereby promoting mild-PTT.

The JC-1 assay was performed to determine the mitochondrial membrane potential, which further confirmed that the combination of nanozyme-catalyzed therapy and mild-PTT could disrupt mitochondria. As shown in Figure 4g, the control group exhibited bright red fluorescence, indicating healthy mitochondria, while the PtNiBiSnSb + NIR group showed very little red fluorescence, with the mitochondrial membrane potential depolarized and manifested by green monomer fluorescence. Cell apoptosis analysis was performed using the Annexin V-FITC/PI apoptosis detection kit. The PtNiBiSnSb + NIR group efficiently induced cell apoptosis due to the synergistic effects of nanozyme-catalyzed reactions and mild-PTT.

It has been reported that both oxidative stress and PTT can induce immunogenic cell death (ICD). Therefore, three ICD markers (ATP, Calreticulin (CRT), and high mobility group protein B1 (HMGB1) were detected to verify whether PtNiBiSnSb could induce ICD. ATP, as a "find-me" signal, is an important chemotactic factor for recruiting immune effector cells. The release of ATP was first measured. The amount of ATP released by PtNiBiSnSb was 10.2 times that of the control group, while the PtNiBiSnSb+NIR group released 15.7 times more ATP than the control group. Immunofluorescence staining experiments revealed that both PtNiBiSnSb and PtNiBiSnSb+NIR groups could induce CRT exposure. The PtNiBiSnSb+NIR group showed brighter green fluorescence, indicating a stronger "eat-me" signal, which was beneficial for enhancing the uptake of tumor cells and cell debris by antigen-presenting cells. Conversely, the PtNiBiSnSb+NIR group exhibited almost no HMGB1 fluorescence, suggesting that it promotes HMGB1 release. HMGB1 facilitates the stable binding of dendritic cells (DCs) to dying tumor cells. Although cell experiments have validated that PtNiBiSnSb can induce ICD, the TME contains many immunosuppressive cells that impede the immune response. These immunosuppressive cells uptake lipids through CD36 as their primary energy source. Therefore, anti-CD36 antibodies were conjugated to the surface of PtNiBiSnSb to block lipid uptake, thereby disrupting the immunosuppressive TME. The ability of PtNiBiSnSb-anti-CD36 to inhibit M2-type macrophages was verified using RAW264.7 cells. To simulate a lipid-rich TME, free fatty acids (palmitic acid) were added to the culture medium. As shown in the Figure 4h, the M2 macrophages (CD 206 as M2 macrophage marker) in the PtNiBiSnSb-anti-CD36+L group decreased by 24.2%, whereas the M2 macrophages in the PtNiBiSnSb+L group decreased by only 17.6%. This indicates that the PtNiBiSnSb-anti-CD36+L group not only induces ICD but also inhibits M2 macrophages by blocking fatty acid uptake. Afterwards, the maturation of DCs was examined. As shown in the Figure 4i, in the PtNiBiSnSb-anti-CD36+L group, the proportion of mature DCs increased by 40.3% compared to the control group. These experimental results manifest that PtNiBiSnSb-anti-CD36+L can effectively promote immune responses. Eventually, a scratch assay was conducted to determine whether the PtNiBiSnSb-anti-CD36 could inhibit cancer cell migration. Compared to the control group, where most cells migrated, the PtNiBiSnSb-anti-CD36+L group showed almost no cell migration (Figure 4j). Collectively, PtNiBiSnSb-anti-CD36 can enhance the immune response and inhibit cancer cell migration (Figure 4k).

### 2.4 Evaluation of in Vivo Immune Response Promotion

Encouraged by the promising results of immune response enhancement in vitro, whether PtNiBiSnSb-anti-CD36 can inhibit immunosuppressive cells was tested in vivo (**Figure 5a**). First, Treg T cells in the spleen (Figure 5b) and tumor were investigated. Compared to the PtNiBiSnSb group, the PtNiBiSnSb-anti-CD36 group indicated a stronger ability to inhibit Treg T cells, while the PtNiBiSnSb-anti-CD36+1064 nm group exhibited even more efficient inhibition of Treg T cells. Then, the macrophages in the tumor tissue were then evaluated (Figure 5c). As expected, PtNiBiSnSb-anti-CD36 reduced the proportion of M2 macrophages (CD 206 as M2 macrophage marker). Compared to the control group, the PtNiBiSnSb-anti-CD36 group presented a decrease of 7.43% (from 11.1% to 3.67%), and the PtNiBiSnSb-anti-CD36+1064 nm group suggested a decrease of 10.39% (from 11.1% to 0.71%). A significant increase in the proportion of M1 macrophages (CD 86 as M1 macrophage marker) was observed in the PtNiBiSnSb-anti-CD36+1064 nm group, rising from 1.01% in the control group to 13.5%. This demonstrates that PtNiBiSnSb-anti-CD36 can inhibit the function of immunosuppressive cells and reverse the immunosuppressive TME. Thereafter, the ability of PtNiBiSnSb-anti-CD36 to enhance the immune response was explored. First, its ability to stimulate DC maturation in vivo was verified. As shown in Figure 5d, the maturation rate of tumor-infiltrating lymph node DCs significantly increased in the PtNiBiSnSb-anti-CD36 group (from 11.5% to 26.6%), while the PtNiBiSnSb-anti-CD36+1064 nm treatment group demonstrated an increase from 11.5% to 47.1%. Then, T cell activation was evaluated. Similar to DC cells, the number of CD4+ T cells and CD8+ T cells in the PtNiBiSnSb-anti-CD36 treatment group increased by 13.2% and 7.4%, respectively, compared to the control group (Figure 5e). All experimental results confirm that PtNiBiSnSb-anti-CD36-mediated ICD, along with interference in lipid metabolism, can reverse the tumor immune microenvironment and promote an anti-tumor immune response.

**Figure 5.**
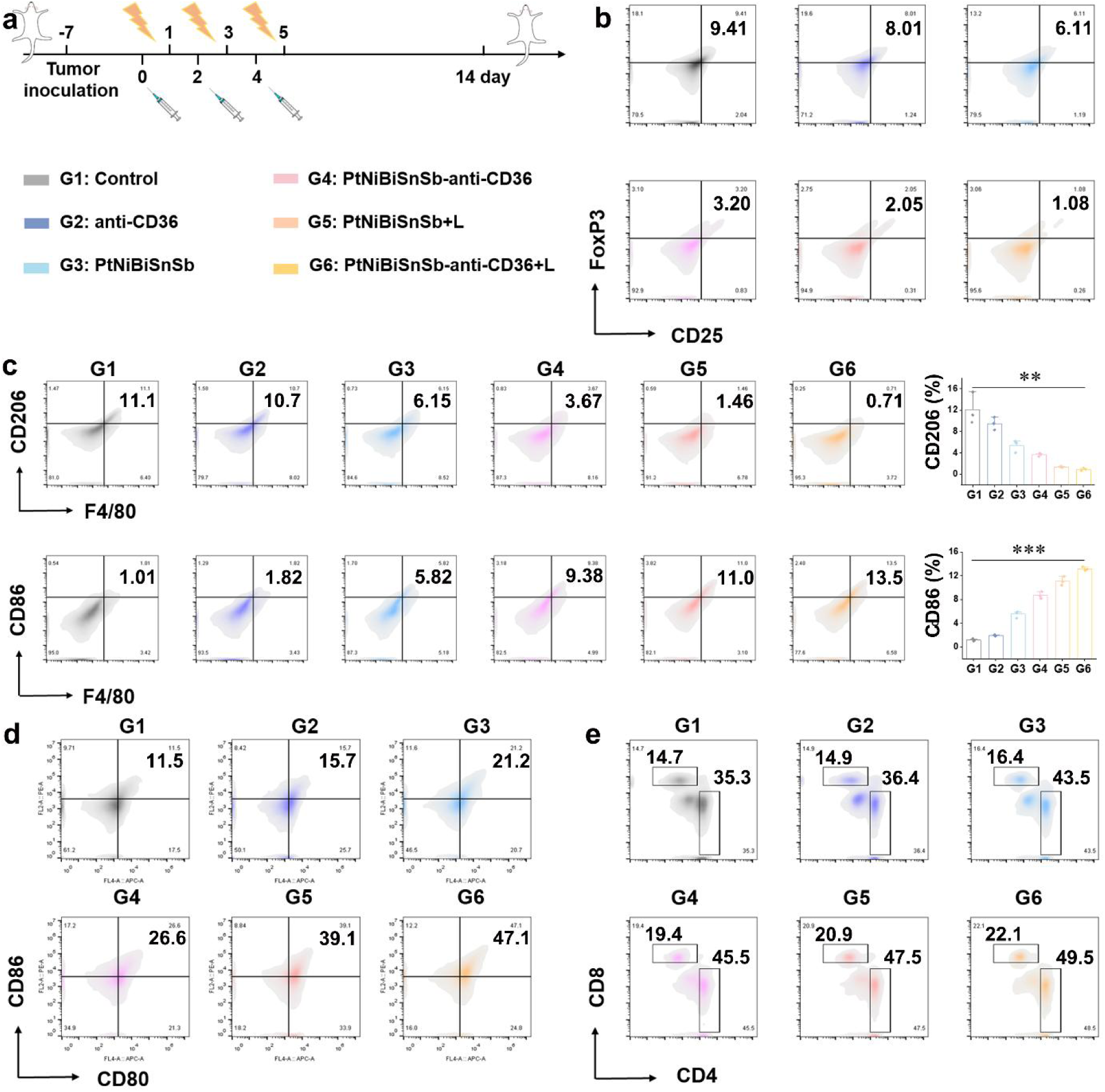
Evaluation of in vivo immune response promotion. (a) Schematic establishment and treatment of the 4T1 tumor-bearing mouse models. (b) Flow cytometry analysis of Treg T cells in spleen after different treatments. (c) CD206 and CD86 expression determined by flow cytometry. Flow cytometric analyses of the populations of (d) DC cells, (e) CD4^+^ T cells, and CD8^+^ T cells. G1: Control, G2: anti-CD36, G3: PtNiBiSnSb, G4: PtNiBiSnSb-anti-CD36, G5: PtNiBiSnSb+1064 nm, G6: PtNiBiSnSb-anti-CD36+1064 nm. *p < 0.05, **p < 0.01, and ***p < 0.001 by Student’s two-tailed t test.

### 2.5. In vivo Anti-tumor Therapy and Anti-metastasis

Because PtNiBiSnSb-anti-CD36 can elicit a strong anti-tumor immune response, its anti-tumor therapeutic effect in vivo was subsequently studied. Before treatment, in vivo fluorescence imaging was performed on the mice. ICG-labeled PtNiBiSnSb was intravenously injected, and the fluorescence intensity in the mice was monitored at 2, 6, 12, 24, 48, and 72 h. As shown in **Figure 6a**, the fluorescence intensity of the PtNiBiSnSb in the tumor was the strongest at 24 h, and the blood circulation half-life was calculated to be τ_1/2(α)_ = 0.98 h and τ_1/2(β)_ = 7.50 h (Figure 6b). The moderate blood circulation time is beneficial for the accumulation of the PtNiBiSnSb at the tumor site. To further verify that the PtNiBiSnSb could target tumor tissue, an in vivo distribution test was conducted. After intravenous injection of PtNiBiSnSb-anti-CD36, the Pt content in major organs (heart, liver, spleen, lung, kidney, tumor) was measured at different time points. As shown in Figure 6c, the content of Pt in the tumor was highest at 24 h post-injection, which is attributed to the EPR (enhanced permeability and retention) effect of the tumor tissue. In addition, the strong fluorescence observed in the kidneys, along with the detection of Pt in mouse feces, suggests that PtNiBiSnSb-anti-CD36 may be excreted through both renal and hepatobiliary pathways. Thereafter, the mice were exposed to a 1064 nm laser 24 h after the injection of PtNiBiSnSb-anti-CD36. The temperature gradually increased with prolonged irradiation, indicating that the PtNiBiSnSb could accumulate in the tumor site 24 h after injection and could be used for mild-PTT.

**Figure 6.**
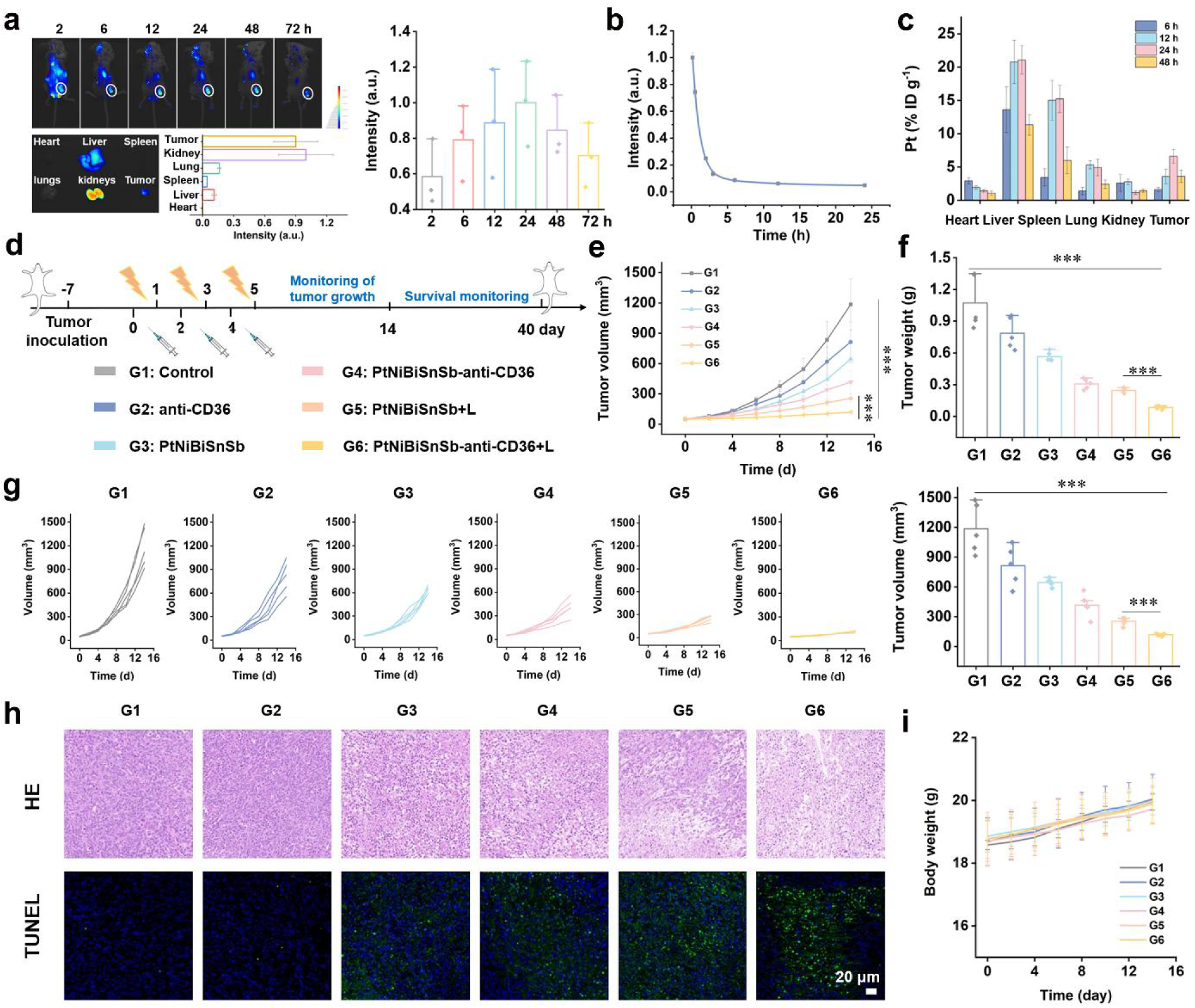
Evaluation of in vivo anti-tumor therapy. (a) In vivo fluorescence imaging after intravenous treatment with PtNiBiSnSb-ICG. Relative quantification of tumoral fluorescence and ex vivo fluorescence images of the major organs and tumor at 72 h post injection of PtNiBiSnSb-ICG. (n=3) (b) Blood circulation after the injection of PtNiBiSnSb-ICG intravenously. (c) Bio-distribution of Pt in major organs and tumors of mice after injection PtNiBiSnSb-anti-CD36 at different time points. (d) Schematic establishment and treatment of the 4T1 tumor-bearing mouse models. (f) Tumor weight, (e, g) tumor growth curves, (h) H&E staining, tunel staining (blue: cell nucleus, green: tunel) were performed on tumor tissues after different treatments. (i) Change curves for the body weight of mice (days 1–14). G1: Control, G2: anti-CD36, G3: PtNiBiSnSb, G4: PtNiBiSnSb-anti-CD36, G5: PtNiBiSnSb+1064 nm, G6: PtNiBiSnSb-anti-CD36+1064 nm.*p < 0.05, **p < 0.01, and ***p < 0.001 by Student’s two-tailed t test.

The in vivo anti-tumor treatment experiment was conducted according to the protocol shown in Figure 6d. As shown in Figures 6e-g, after 14 days of treatment, compared to the control group and the CD36 antibody alone group, the PtNiBiSnSb group could moderately inhibit the growth of tumors due to the generation of large amounts of ROS in the TME, leading to apoptosis. PtNiBiSnSb-anti-CD36 showed better therapeutic effects than pure PtNiBiSnSb because it interfered with lipid metabolism. The most effective treatment was observed in the PtNiBiSnSb-anti-CD36+1064 nm group, with inhibition rates of 92.1% for tumors. To further confirm its anti-tumor effects, histological and apoptotic studies were performed on tumor tissue sections using H&E, TUNEL, and Ki67 staining. As shown in Figure 6h, compared to the control group, the tumor tissues in the PtNiBiSnSb-anti-CD36+1064 nm group were severely damaged. Furthermore, the TUNEL fluorescence was the brightest in this group, implying the highest number of apoptotic cells. Ki-67, a protein highly expressed during cell division, was effectively inhibited in the PtNiBiSnSb-anti-CD36+1064 nm group. Therefore, the combination of nanozyme-catalyzed therapy, mild-PTT, and lipid metabolism interference can effectively activate immune responses and enhance anti-tumor immunotherapy. Besides, there were no significant fluctuations in the body weight of mice in different groups during the entire treatment period (Figure 6i). H&E-stained sections validated that the tissue structure of major organs such as the heart, liver, spleen, lungs, and kidneys remained intact, and blood biochemical indicators were normal, indicating that the biosafety of PtNiBiSnSb-anti-CD36.

Currently, 90% of cancer-related deaths are due to cancer metastasis and recurrence. A material that can inhibit both tumor growth and metastasis and recurrence would be highly valuable for clinical applications. The ability of PtNiBiSnSb-anti-CD36 to inhibit metastasis was evaluated through a lung metastasis experiment. As shown in **Figure 7a**, numerous tumor nodules were noticed in the control group. In the PtNiBiSnSb-anti-CD36 group, the number of tumor nodules was significantly reduced, and in the PtNiBiSnSb-anti-CD36+L group, no tumor nodules were noted, signifying that PtNiBiSnSb-anti-CD36+L effectively inhibits lung metastasis. Next, we tested whether the tumors in the mice still exhibited an immune response on the 26th day. First, immunofluorescence staining of the tumor tissue was performed to test if PtNiBiSnSb-anti-CD36+L treatment could induce significant CRT exposure. The experimental results showed that a considerable amount of CRT fluorescence could still be observed in the PtNiBiSnSb-anti-CD36+L group (Figure 7b). Then, tumor tissue immunofluorescence staining (Figure 7c) and flow cytometry (Figure 7d) were conducted to examine CD4/CD8 T cell infiltration in the tumor cells. The results from both experiments were consistent, indicating significant CD4/CD8 T cell infiltration in the tumor tissue after PtNiBiSnSb-anti-CD36+L treatment. Subsequently, memory T cells were evaluated (Figure 7e), showing that the proportion of memory T cells in the PtNiBiSnSb-anti-CD36+L group was significantly higher than in the control group. Due to the synergistic effect of PtNiBiSnSb-anti-CD36 treatment, the survival rate of tumor-bearing mice was significantly increased (Figure 7f). Furthermore, the enhanced secretion of inflammatory cytokines TNF-α (Figure 7g) and IFN-γ (Figure 7h) further indicated that PtNiBiSnSb-anti-CD36 treatment effectively activated the anti-tumor immune response in mice. Therefore, nanozyme-catalyzed therapy, combined with mild-PTT and interference with lipid metabolism, can effectively prevent tumor recurrence and metastasis.

**Figure 7.**
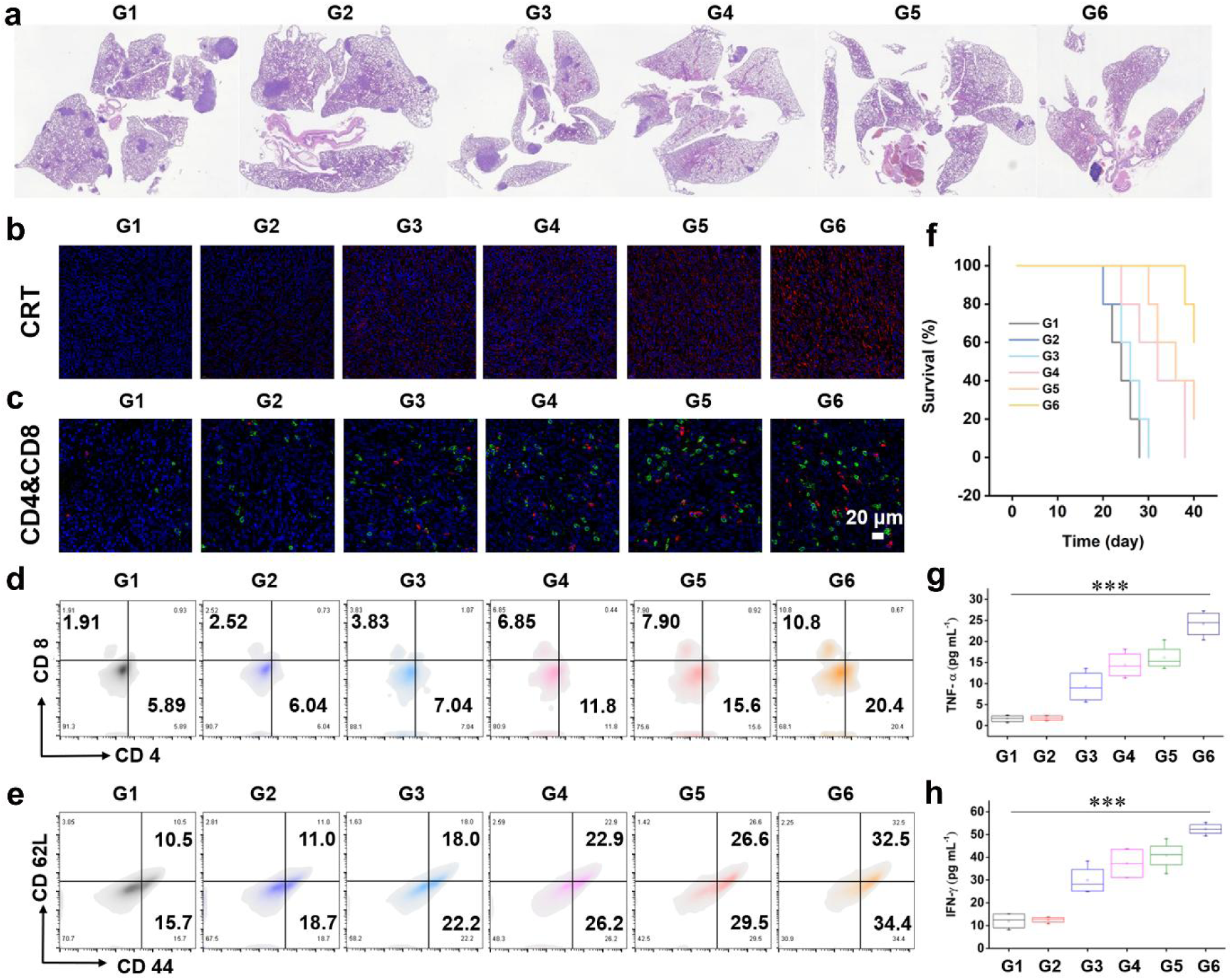
Evaluation of in vivo anti-metastasis. (a) H&E staining of lung tissues with different treatments. (b) CRT staining (blue: cell nucleus, red: CRT), (c) CD4&CD8 staining (blue: cell nucleus, red: CD4, green: CD8) were performed on tumor tissues after different treatments. (d) Flow cytometric analyses of the populations of CD4^+^ T cells and CD8^+^ T cells in tumor. (e) Flow cytometric analyses of the populations of memory T cells. (f) Survival rate of mice after different treatments. (g,h) Pro-inflammatory cytokine (TNF-α and IFN-γ) secretion upon different treatments. *p < 0.05, **p < 0.01, and ***p < 0.001 by Student’s two-tailed t test.

## 3. Conclusion

In this paper, a PtNiBiSnSb-anti-CD36 HEAzyme was constructed to promote cancer immunotherapy by boosting enzyme-like catalytic ability and interfering with dual energy metabolism. The PtNiBiSnSb-anti-CD36 HEAzyme exhibits both POD-like and NADH-POD-like activities. With its abundant active sites, the HEAzyme demonstrates superior catalytic activity compared to SAzymes. Kinetic studies demonstrate that the catalytic activity of PtNiBiSnSb HEAzyme is 7.2 times greater than that of its structurally analogous but non-HEAzyme counterpart (PtBi). DFT calculations and machine learning predictions provide a deep understanding of the enhanced POD-like activity mechanism. With the addition of Ni, Sn, and Sb, the d-band center of Pt sites on the surface of HEAzyme gradually decreases, suppressing the excessive binding of hydroxyl groups on the HEAzyme surface and enhancing the POD like properties of the PtNiBiSnSb. Additionally, the synergistic effects of enzymatic therapy and mild-PTT induce ICD. The enhanced POD-like activity generates significant amounts of ROS and depletes NADH within the TME, disrupting normal energy metabolism. Furthermore, PtNiBiSnSb-anti-CD36 disrupts lipid metabolism by inhibiting fatty acid uptake, leading to effective tumor growth suppression. The interference with lipid metabolism also impairs the function of immunosuppressive cells, further promoting the immune response alongside ICD. Both in vitro and in vivo experiments confirm that PtNiBiSnSb-anti-CD36 effectively induces ICD and achieves the goals of inhibiting immunosuppressive cells, tumor cells, and cancer cell metastasis. This work not only establishes a precedent for using HEAzyme in cancer immunotherapy but also deepens the theoretical understanding of HEAzyme’s role in POD-like reactions.

## Conflict of Interest

The authors declare no conflict of interest.

Received: ((will be filled in by the editorial staff))

Revised: ((will be filled in by the editorial staff))

Published online: ((will be filled in by the editorial staff))

---

**Scheme 1.**
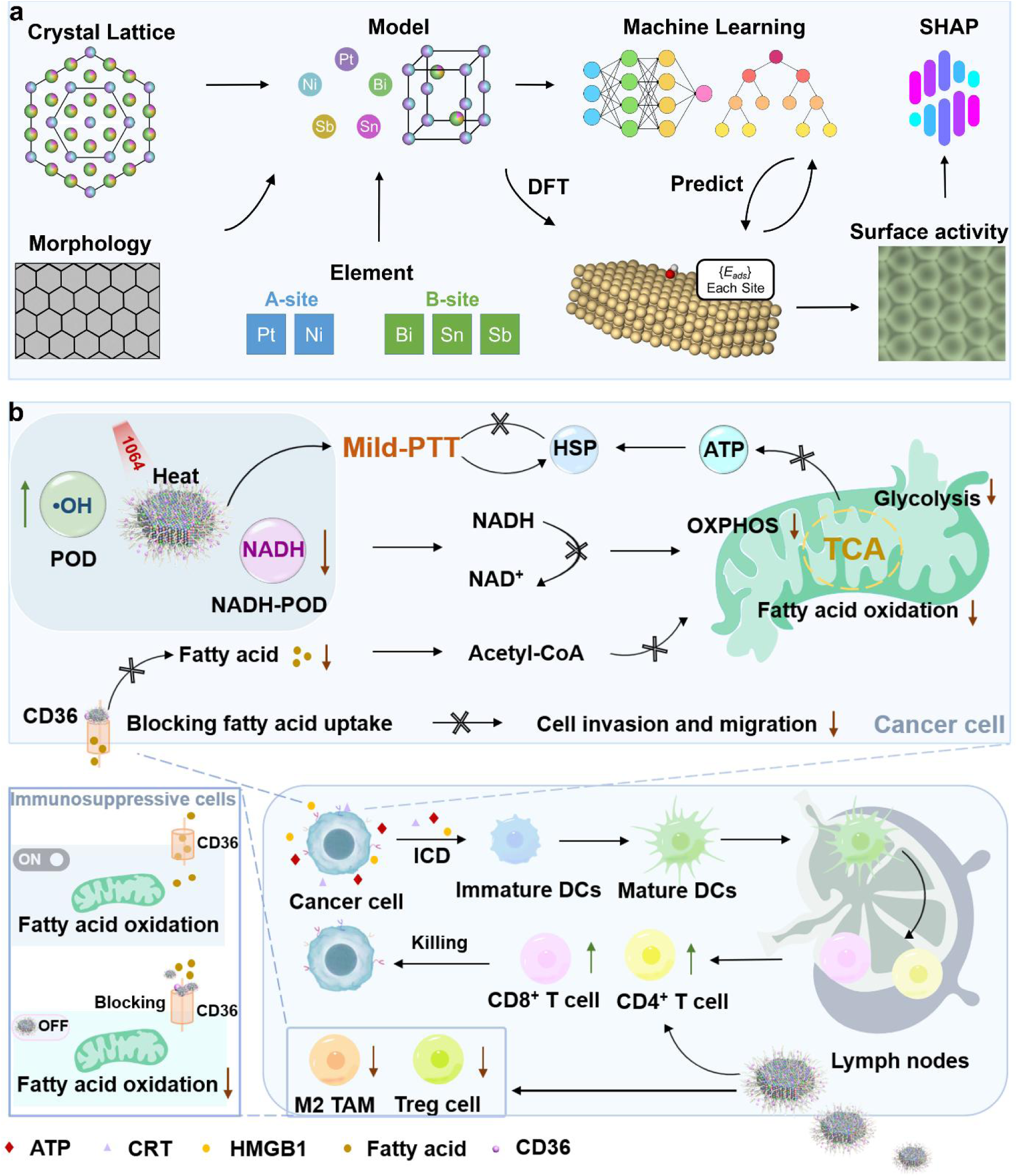
Schematic illustration of machine-learning-assisted PtNiBiSnSb-anti-CD36 for cancer immunotherapy. (a) Machine-learning assisted DFT process and (b) PtNiBiSnSb-anti-CD36 for cancer immunotherapy process diagram.

A high-entropy atom nanozyme (HEAzyme) PtNiBiSnSb-anti-CD36 with enhanced POD-like and NADH-POD activities was designed for efficient anti-tumor immunotherapy. The mechanism of enhanced POD-like enzyme activity in PtNiBiSnSb was explained through machine learning predictions and DFT calculations. This PtNiBiSnSb-anti-CD36 HEAzyme not only achieves enhanced oxidative stress within the tumor microenvironment and interferes with energy metabolism to inhibit cancer cell growth, but also promotes cancer immunotherapy by inhibiting lipid metabolism.

*Man Wang, Wen Chen, Zhicheng Zhu, Yulin Xie, Guoqing Zhu, Yanrong Qian, Mengyu Chang, Shuiping Luo* Zhiling Zhu,* and Chunxia Li\**

TOC figure

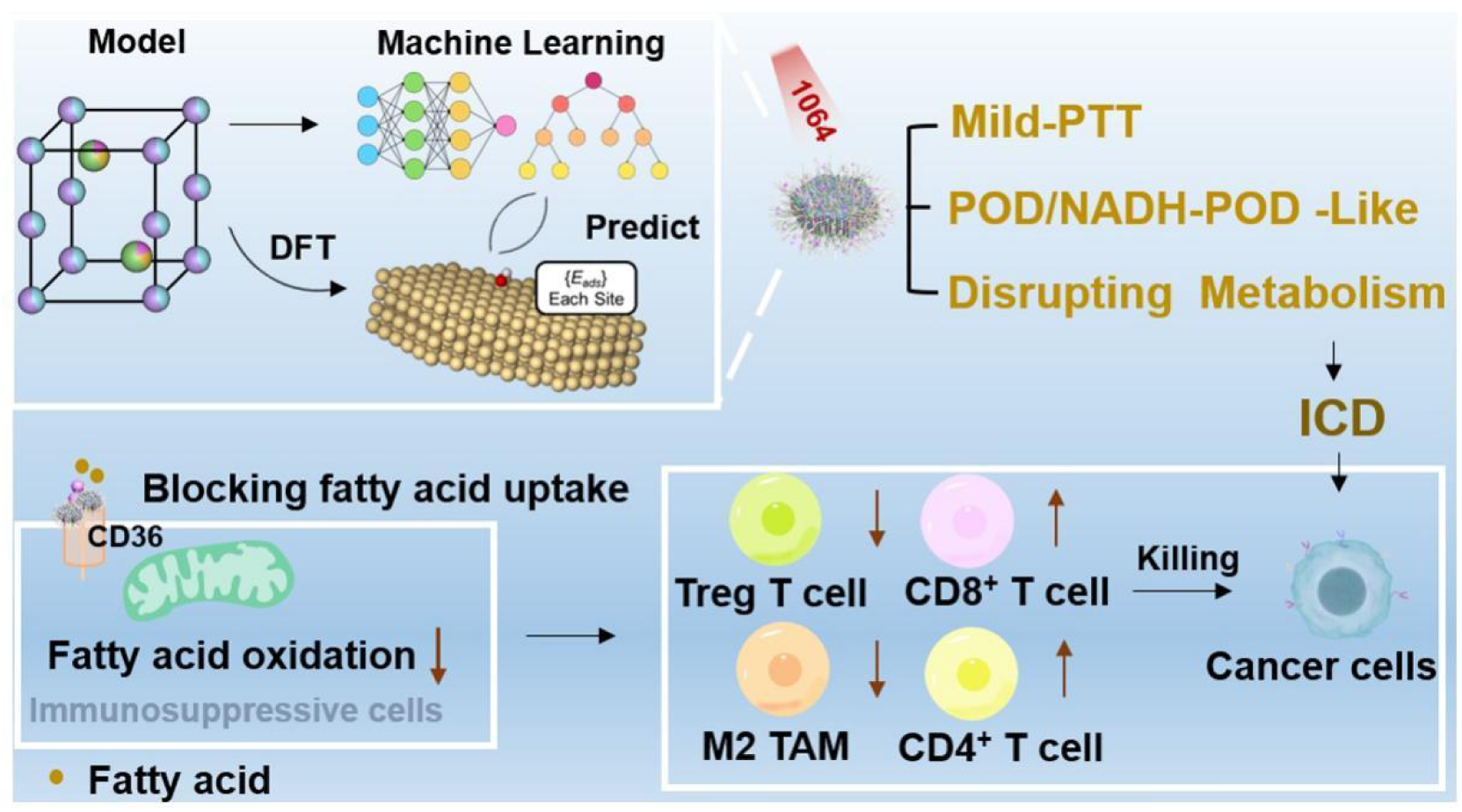

## Notes

### Competing Interest Statement

The authors have declared no competing interest.

